# Molecules inhibit the enzyme activity of 3-chymotrypsin-like cysteine protease of SARS-CoV-2 virus: the experimental and theory studies

**DOI:** 10.1101/2020.05.28.120642

**Authors:** Zhesheng He, Wencong Zhao, Wenchao Niu, Xuejiao Gao, Xingfa Gao, Yong Gong, Xueyun Gao

## Abstract

SARS-CoV-2 has emerged as a world public health threat. Herein, we report that the clinical approved auranofin could perfectly inhibit the activity of 3-chymotrypsin-like cysteine protease (Mpro or 3CLpro) of SARS-CoV-2. Gold cluster could significantly inhibit 3CLpro of SARS-COV-2. Phenyl isothiocyanate and Vitamin K3 could well suppress the activity of 3CLpro. For Mpro inhibition, IC50 of auranofin, Vitamin K3, phenyl isothiocyanate, gold cluster are about 0.51μM, 7.96μM, 10.13μM, 1.61μM, respectively. These compounds may be with potentials for treatment SARS-CoV-2 virus replication. Especially for FDA approved auranofin, it is an anti-inflammation drug in clinic, thus it may with strong potential to inhibit virus replication and suppress the inflammation damage in COVID-19 patients. Gold cluster is with better safety index and well anti-inflammation in vitro/vivo, therefore it is with potential to inhibit virus replication and suppress the inflammation damage caused by COVID-19 virus. As Au(I) ion is active metabolism specie derived from gold compounds or gold clusters in vivo, further computational studies revealed Au ion could tightly bind thiol group of Cys145 residue of 3CLpro thus inhibit enzyme activity. Also, phenyl isothiocyanate and Vitamin K3 may interact with thiol group of Cys145 via Michael addition reaction, molecular dynamic (MD) theory studied are applied to confirmed these small molecules are stable in the pocket and inhibit Mpro activity.

## Introduction

A new coronavirus named as COVID-19 virus (also called SARS-CoV-2 virus) is responsible for the 2020 pandemic outbreak in the world. The viral 3-chymotrypsin-like cysteine protease (3CLpro, also called Mpro) enzyme controls this COVID-19 virus replication and is essential for its life cycle. Therefore, 3CLpro is drug target in the case of SARS-CoV-2. Our biochemistry studied revealed that the gold compounds such as auranofin, Isothiocyanate compounds such as phenyl isothiocyanate, vitamin K such as vitamin K3, and gold cluster such as glutathione coated gold cluster could well inhibit the activity of 3CLpro. Further DFT and MD computational studies also verified these small molecules could interact Cys145 and amino acid residues of Mpro, thus suppress the activity of Mpro. These compounds could serve as potential anti-SARS-CoV-2 lead molecules for further drug studies to combat COVID-19.

### Expression and purification of 3CLpro

Recombinant COVID-19 virus 3CLpro with native N and C termini was expressed in Escherichia coli and subsequently purified following the recently reported work (1). The full-length gene encoding COVID-19 virus Mpro was synthesized for Escherichia coli expression. Briefly, the expression plasmid was transformed into Escherichia coli cells and cultured in Luria Broth medium containing 100 μg/ml ampicillin at 37 °C. When the cells were grown to OD600 of 0.6-0.8, 0.5 mM IPTG was added to the cell culture to induce the expression at 16 °C. After 10 h, the cells were harvested by centrifugation at 3,000g. The cell pellets were resuspended in lysis buffer (20 mM Tris-HCl pH 8.0, 300 mM NaCl), lysed by high-pressure homogenization, and then centrifuged at 25,000g for 40 min. The supernatant was loaded onto Ni-NTA affinity column, and washed by the resuspension buffer containing 20 mM imidazole. The His tagged Mpro was eluted by cleavage buffer (50 mM Tris-HCl pH 7.0, 150 mM NaCl) including 300 mM imidazole. Human rhinovirus 3C protease was added to remove the C-terminal His tag. The Mpro was further purified by ion exchange chromatography and size exclusion chromatography. The purified Mpro was stored in 50 mM Tris-HCl pH 7.3, 1 mM EDTA.

### Inhibit Enzyme activity of Mpro in vitro

In order to characterize 3CLpro enzymatic activity, we used a fluorescence resonance energy transfer (FRET) assay (1, 2). To do this, a fluorescently labeled substrate, (EDNAS-Glu)-Ser-Ala-Thr-Leu-Gln-Ser-Gly-Leu-Ala-(Lys-DABCYL)-Ser, derived from the auto-cleavage sequence of the viral protease was chemically modified for enzyme activity assay (2). Potential inhibitors suppress COVID-19 virus Mpro were screened by this enzymatic inhibition assay. Molecules picked from gold compounds library, gold cluster library (gold atoms stack together to form cluster and the cluster is stabled by organic molecule with thiol group, where thiol groups interact with gold cluster via Au-S bond), Isothiocyanate library, and vitamin K library were used. When the different compounds were added into the enzymatic reaction solution, the change of initial rates was calculated to evaluate their inhibitory effect. The molecule inhibit Mpro activity reaction mixture included 0.5 μM protein, 20 μM substrate (EDNAS-Glu)-Ser-Ala-Thr-Leu-Gln-Ser-Gly-Leu-Ala-(Lys-DABCYL)-Ser) and molecules from afore mentioned compound library. The compounds of interest were defined as those with a percentage of inhibition over 50% compared with the reaction in the absence of inhibitor. IC50 values of four compounds were measured using 0.5 μM protein, 20 μM substrate and 9 different inhibitor concentrations of compounds. All experiments were performed in triplicate.

## Experimental Results and discussion

For Mpro inhibition, IC50 of auranofin is about 0.51μM, data showed as following Figure 1. This is a very strong suppressor for Mpro. This result is a strong proof to well understand why auranofin could well inhibit virus replication (3). The auranofin was approved by FDA in clinical treatment Rheumatoid Arthritis treatments via suppress the immune cytokine activity (4). As COVID-19 virus could cause immune cytokine storm in patients, recent reports revealed drugs inhibit immune cytokine activity could well recover severe patients (5). The auranofin combined the potential of anti-virus and the proved anti-inflammation together to recover severe COVID-19 infected patients.

**Figure 1.**
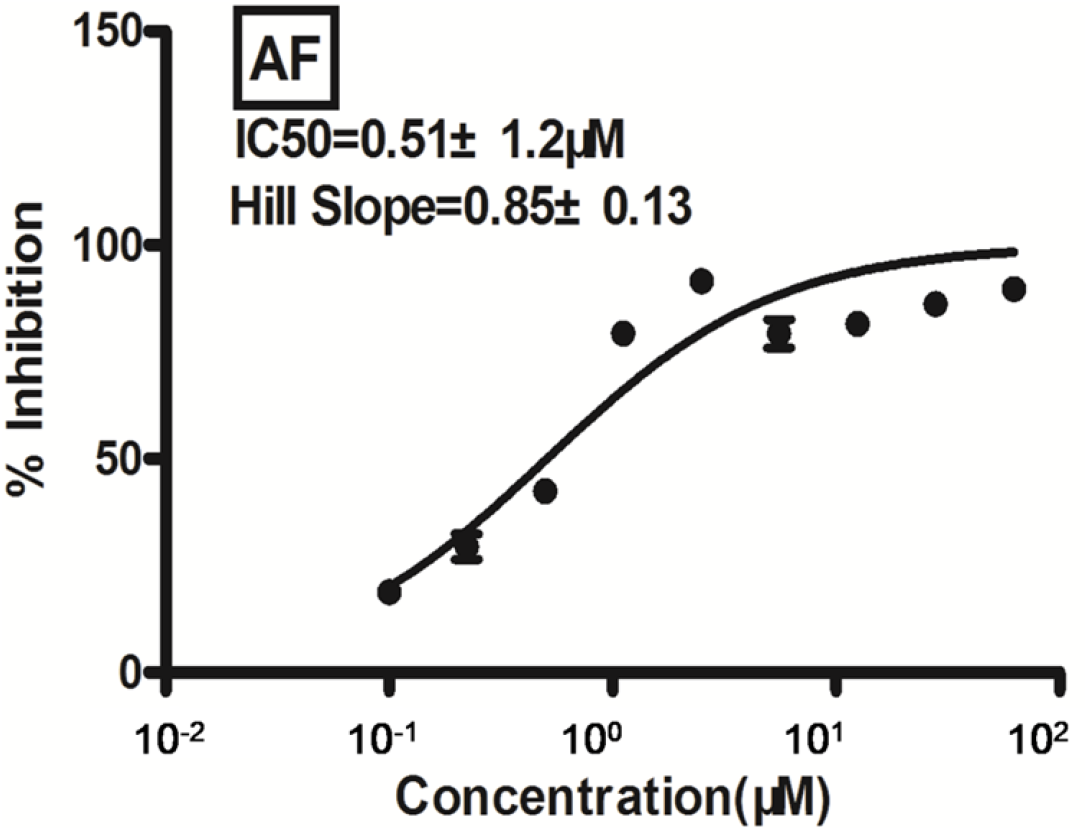
auranofin inhibit Mpro at dose dependent manner.

For Mpro inhibition, IC50 of Menadione (vitamin K3) is about 7.96μM, this is very valuable lead molecule for further anti-virus studies, data showed as following Figure 2. Vitamin K is widely used as dietary supplements approved by FDA (6), thus it is benefit for vitamin K3 applied for further anti-COVID-19 virus studies.

**Figure 2.**
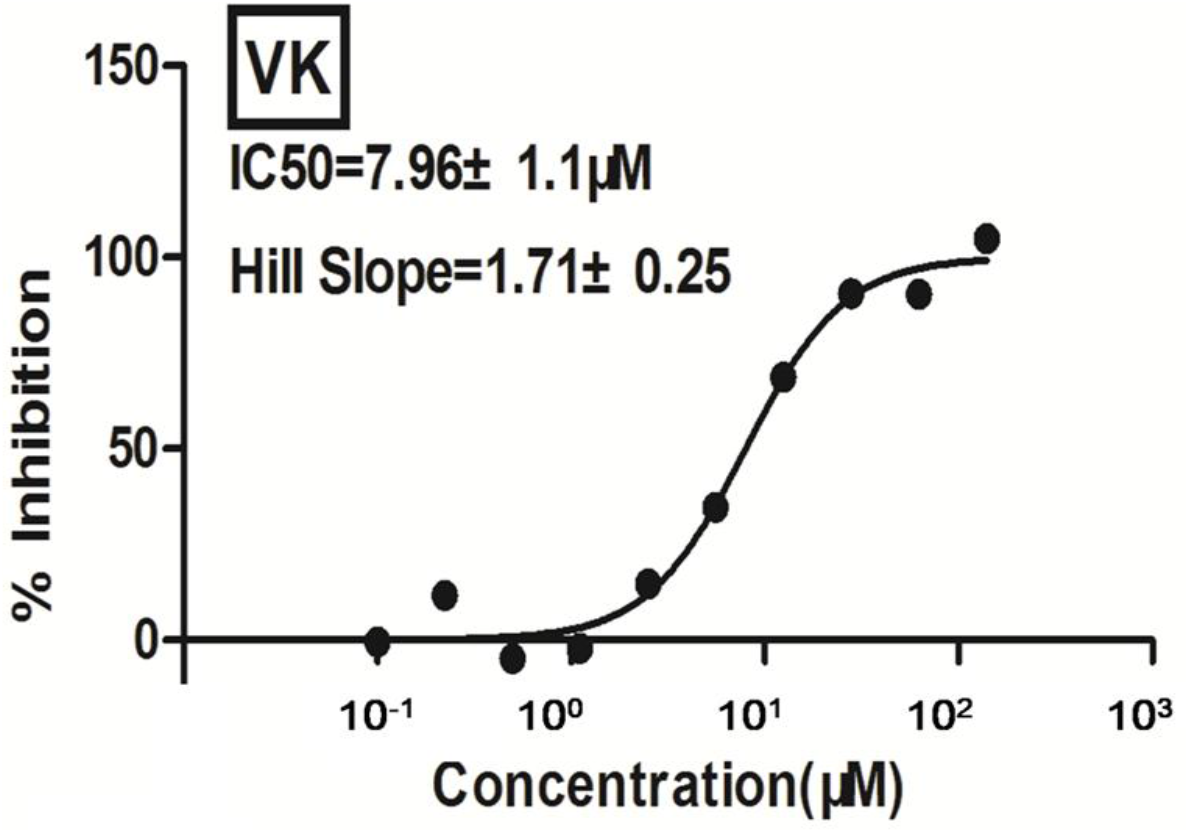
Vitamin K3 inhibit Mpro at dose dependent manner.

For Mpro inhibition, IC50 of phenyl isothiocyanate is about 10.13μM, data showed as following Figure 3. Phenethyl isothiocyanate is a constituent of cruciferous vegetables that has cancer preventive activity including lung, prostate, and breast cancer (7). The potential of this compound to inhibit COUID-19 virus is an interesting topic as phenethyl isothiocyanate is a good candidate to be a dietary supplement.

**Figure 3.**
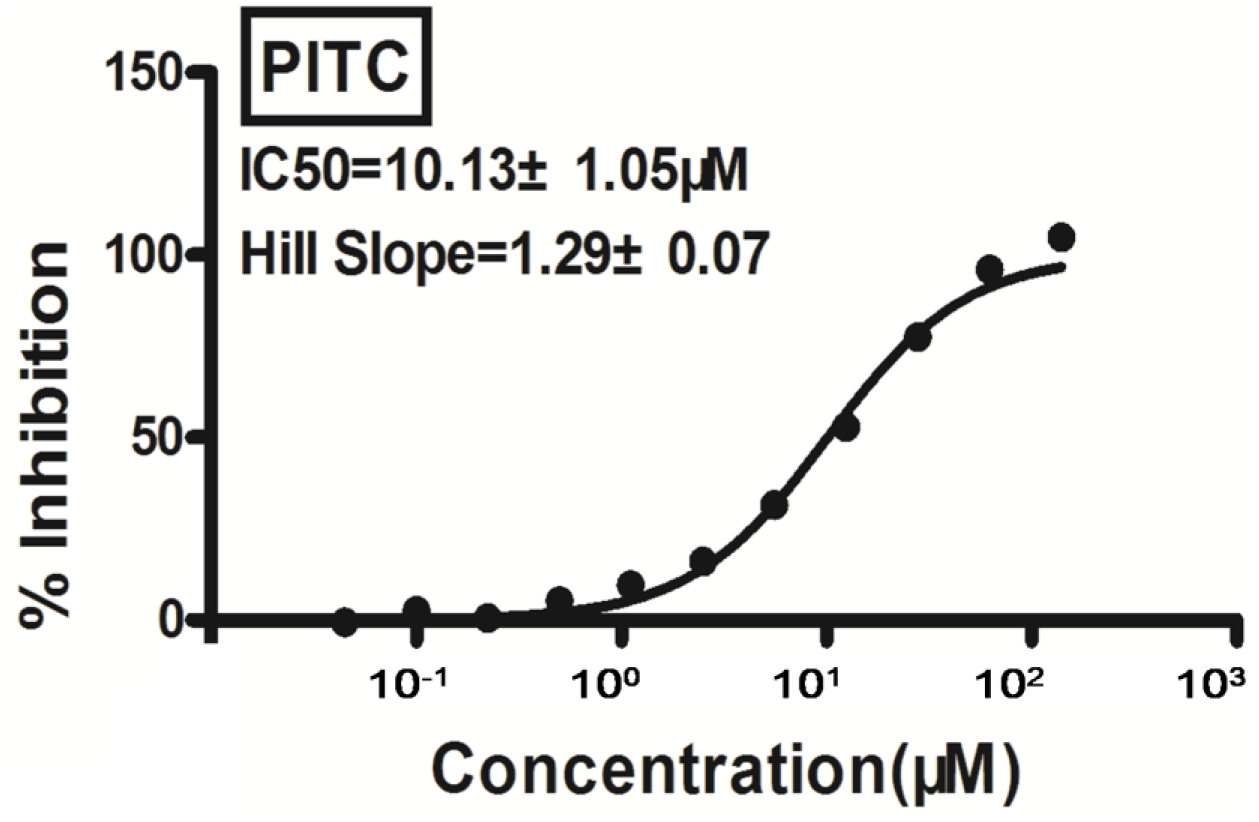
phenethyl isothiocyanate inhibit Mpro at dose dependent manner.

For Mpro inhibition, IC50 of glutathione gold cluster (Au_29_SG_27_) is about 1.61μM, data showed as following Figure 4. This result showed glutathione gold cluster could well inhibit Mpro in vitro. Tt is with strong potentials to inhibit COVID-19 virus replication. Note that gold clusters are with strong anti-inflammation for Rheumatoid Arthritis treatments in vitro/vivo, its safety index is better than FDA approved Auranofin and Methotrexate in animal studies (8, 9, 10, 11). Combined its anti-virus Mpro and clarified anti-inflammation effects, gold cluster may be with advantage in later anti-COVID-19 studies.

**Figure 4.**
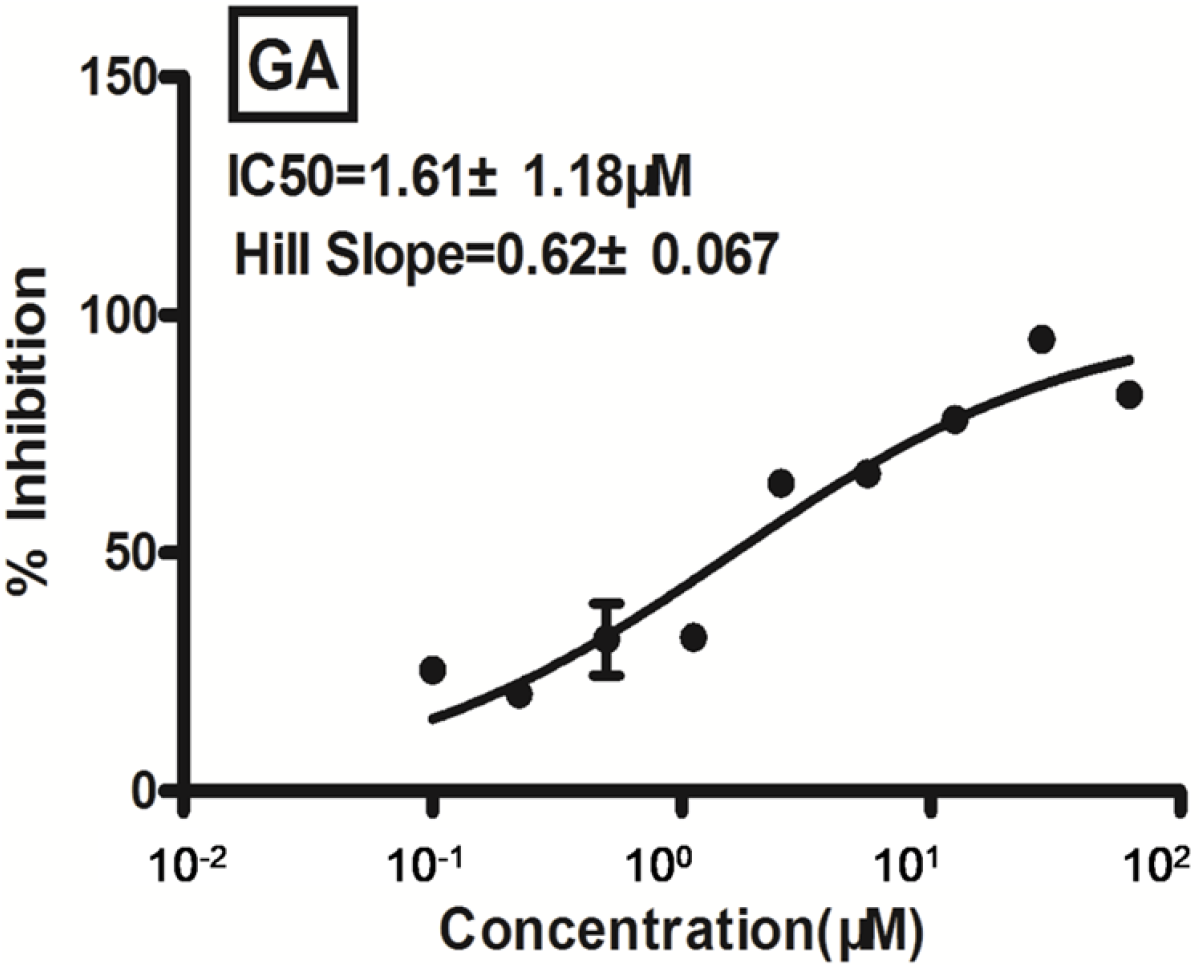
Gold cluster inhibit Mpro at dose dependent manner.

## Theory calculations and discussion

It is widely accepted that gold compounds metabolism in vivo and the produced Au(I) would interact with specific thiol group of protein to interfere their normal functions (12). The interactions between the gold atom with the binding pockets of proteins were studied by density functional theory (DFT) calculations. For the calculations, only the main amino acids of the protein binding pockets were taken into account. All geometries were fully optimized using the B3LYP method in conjunction with the SDD basis set for Au and the 6-31G(d,p) for other nonmetals (13, 14, 15). The SDD pseudopotential was also applied for Au. During optimization, SMD4 solvation model was utilized to model the water environment. All the calculations were carried out using Gaussian 09 package (16). The binding energy (Eb) between Au and the pocket ligands were calculated using the following equation: Eb = EAu+ + Eligands – EAu-ligands. where EAu+, Eligands, and EAu-ligands were the total energies of Au+ ion, the ligands, and the Au-ligand complex structures, respectively. Eligands were the single-point energy for the Au-ligand complex with the Au atom removed from the system.

The interaction between gold atom with the binding pocket of proteins SARS-CoV-2 3CLpro (PDB ID: 6Y2E) was studied by DFT calculations. The binding pocket consisted mainly of a cysteine and a histidine. The Au atom preferred to form a linear S-Au-N bonding configuration with the thiol sulfur atom of Cys145 and the imidazole nitrogen atom of His41. The Au atom had a valence of +1 because of the formation of the Au-S bond. The histidine N atom donated its electron lone pair to the vacant orbital of Au, which did not change the valance of Au. The binding energy was as high as 90.5 kcal mol−1. This suggested that Au+ ion firmly locked the cysteine and histidine groups together, which might hinder the bioactivity of the protein.

For molecular dynamic (MD) simulation, the systems were constructed by SARS-CoV-2 main protease (PDB: 6Y2E) complex with phenyl isothiocyanate or Vitamin K3. The stability of the system was obtained by MD simulation using Nanoscale MD (NAMD) software (18). These input files were generated by CHARMM-GUI (19). All the systems were solvated using the TIP3 water model and neutralized with NaCl to an ionic concentration of 0.15 M. The following simulation protocols were applied for each system: 10 ps minimization by steepest descent method; 5 ns equilibration in standard number of particle, volume and temperature (NVT) ensemble; followed by unrestrained 50 ns production MD simulations in standard number of particle, pressure and temperature (NPT) ensemble, at time step and collection interval of 2 fs and 1 ps, respectively. The stability of the systems was examined by computing the backbone root mean-squared deviation (RMSD) of the entire trajectories.

Once the phenyl isothiocyanate interact with thiol group of Cys145 in the binding pocket of SARS-CoV-2 main protease (PDB: 6Y2E), this small molecule may further interact with close amino acid residues. MD result, Figure 6, showed the binding pocket consisted mainly of His-41, His-164, and Met-165. Phenyl isothiocyanate forms hydrogen bond interaction with His-41 and His-164, as hydrogen atom donor. In addition, the structure of phenyl isothiocyanate was very consistent with the structure of the Mpro pocket.

**Figure 5.**
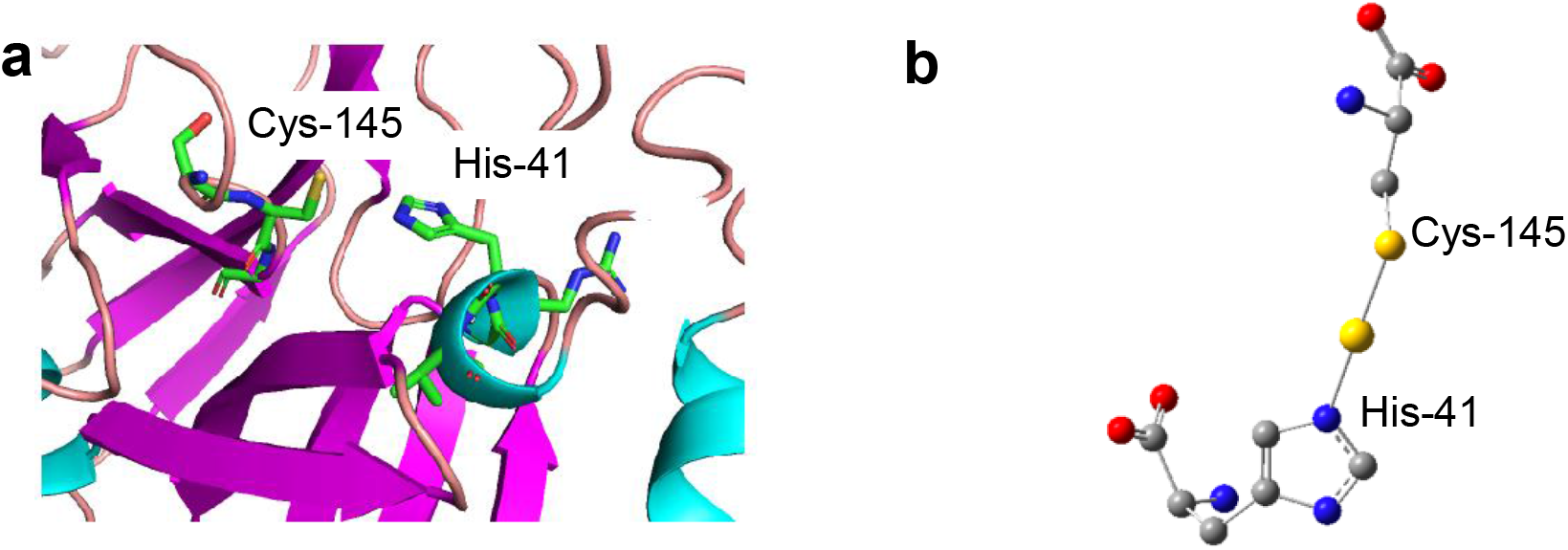
DFT calculation for the coordination interaction between Au+ and the binding pocket of protein SARS-CoV CLpro (PDB ID: 6Y2E). (a) Binding pocket consisting of amino acids Cys-145 and His-41. (b) The stable interaction configuration between Au+ and the amino acids. Hydrogen atoms in b are not shown for clarity.

**Figure 6.**
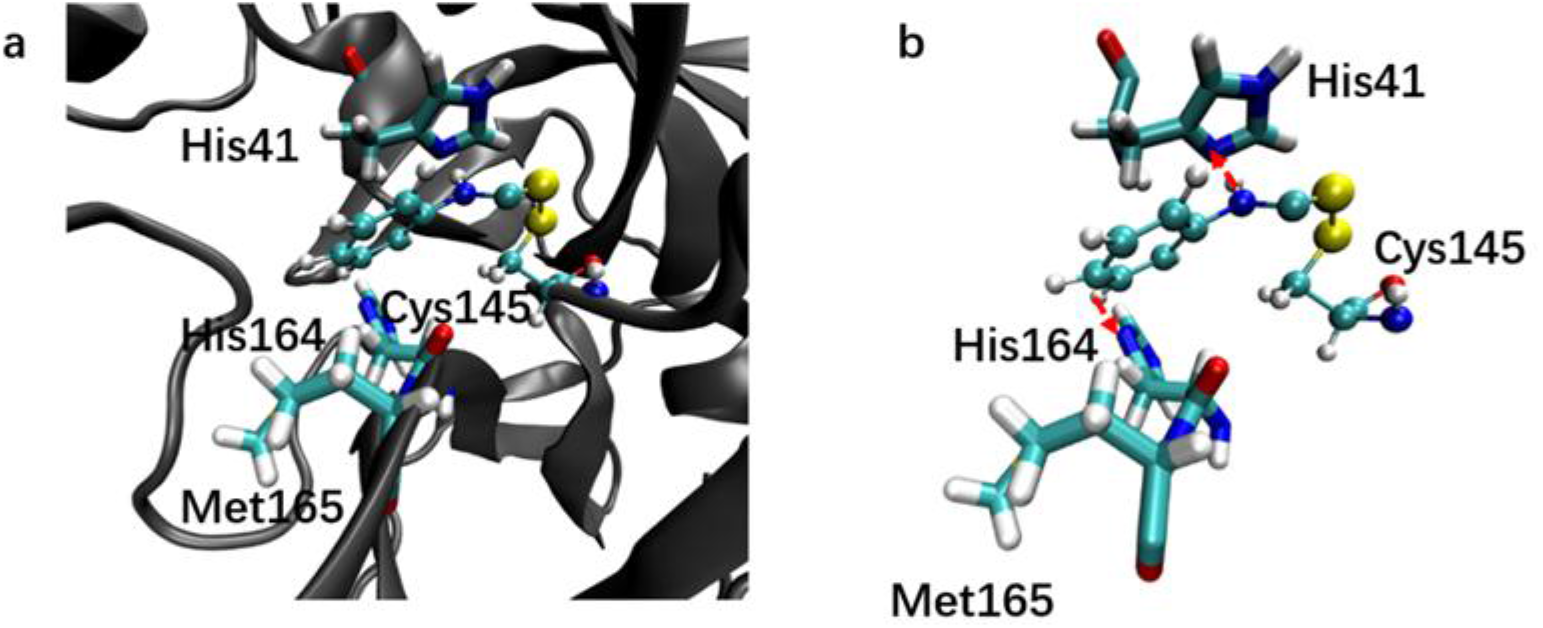
MD simulation for SARS-CoV-2 main protease (PDB: 6Y2E) complex with phenyl isothiocyanate. (a) The binding pocket mainly consisting of amino acids His-41, His-164, and Met-165. (b) The stable interaction configuration between phenyl isothiocyanate and the amino acids. The red arrow indicates the direction of hydrogen bond. Yellow-S, Red-O, Blue-N, Gray-C, White-H.

Similar, the binding pocket consisted mainly of His-41, Gly-143 and Glu-166 of SARS-CoV-2 main protease (PDB: 6Y2E) complex with Vitamin K3, see Figure 7. This small molecule form hydrogen bond interaction with Cly-143 and Glu-165. In addition, the structure of Vitamin K3 was consistent with the shape of the pocket too.

**Figure 7.**
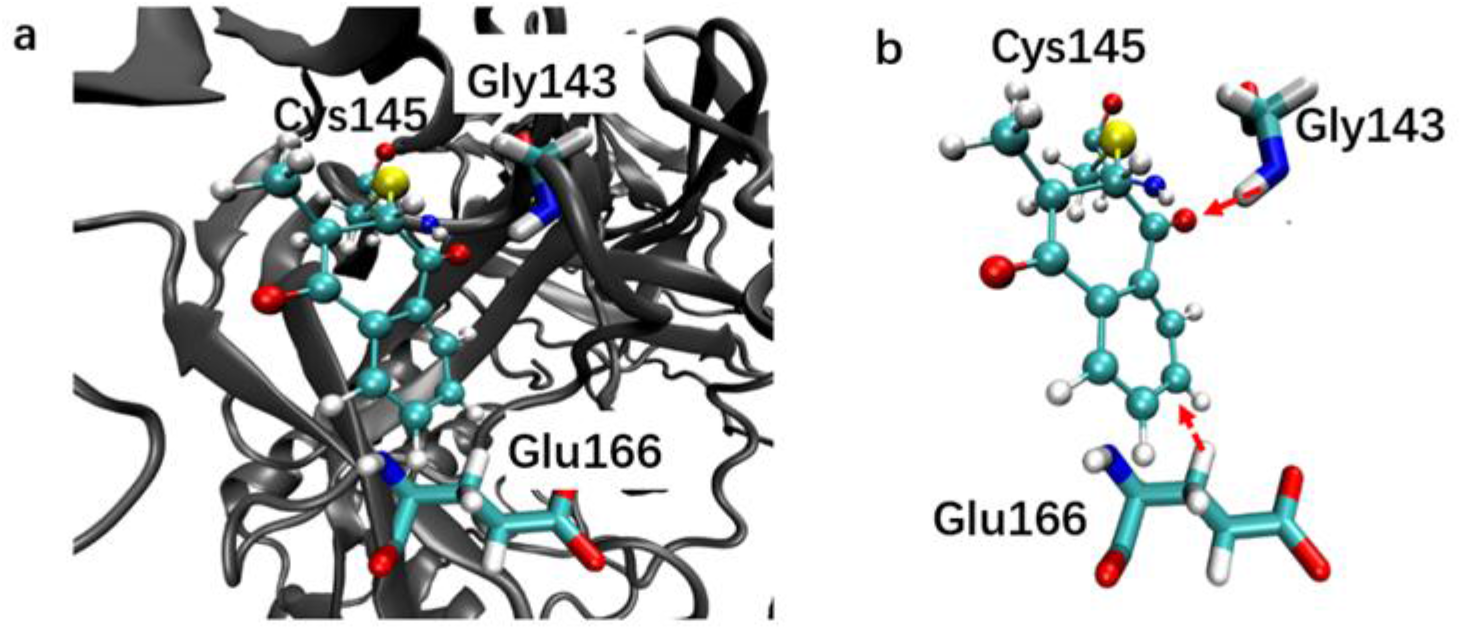
MD simulation for SARS-CoV-2 main protease (PDB: 6Y2E) complex with Vitamin K3. (a) The binding pocket mainly consisting of amino acids Gly-143 and Glu-166. (b) The stable interaction configuration between Vitamin K3 and the amino acids. Yellow-S, Red-O, Blue-N, Gray-C, White-H.

## Notes

### Competing Interest Statement

The authors have declared no competing interest.

### Summary of Updates

Theory studies of the interaction between small molecule and Mpro were carried out. This theory result reveal small molecules could strongly interact with the Cys145 and close amino acid residues, thus suppress the enzyme activity of Mpro.

## Reference

1. Zhenming Jin, Xiaoyu Du, Yechun Xu, Yongqiang Deng, Meiqin Liu, Yao Zhao, Bing Zhang, Xiaofeng Li, Leike Zhang, Chao Peng, Yinkai Duan, Jing Yu, Lin Wang, Kailin Yang, Fengjiang Liu, Rendi Jiang, Xinglou Yang, Tian You, Xiaoce Liu, Xiuna Yang, Fang Bai, Hong Liu, Xiang Liu, Luke W. Guddat, Wenqing Xu, Gengfu Xiao, Chengfeng Qin, Zhengli Shi, Hualiang Jiang, Zihe Rao & Haitao Yang, Nature, 2019, https://doi.org/10.1038/s41586-020-2223-y

2. Valerie Grum-Tokars, Kiira Ratia, Adrian Begaye, Susan C. Baker, Andrew D. Mesecara, Virus Research, 2008, 133(1):63–73

3. Hussin A. Rothan, Shannon Stone, Janhavi Natekar, Pratima Kumari, Komal Arora, Mukesh Kumar, 2020, https://www.biorxiv.org/content/10.1101/2020.04.14.041228v1.full

4. Hull RG, Morgan SH, Parke AL, Childs L, Goldman M, Hughes GR., Int J Clin Pharmacol Res. 1984; 4(6):395–401.

5. Yang Cao., JiaWei., Liang Zou, Tiebin Jiang, Gaoxiang Wang, Liting Chen, Liang Huang, Fankai Meng, Lifang Huang, Na Wang, Xiaoxi Zhou, Hui Luo, Zekai Mao, Xing Chen, Jungang Xie, JingLiu, Hui Cheng, Jianping Zhao, Jianfeng Zhou, Journal of Allergy and Clinical Immunology, 2020, https://doi.org/10.1016/j.jaci.2020.05.019

6. https://ods.od.nih.gov/factsheets/vitaminK-HealthProfessional/

7. Urvi Telang, Marilyn E. Morris, Mol Nutr Food Res. 2010, 54(12): 1802–1806.

8. Fuping Gao, Qing Yuan, Pengju Cai, Liang Gao, Lina Zhao, Meiqing Liu, Yawen Yao, Zhifang Chai, and Xueyun Gao, Adv. Sci., 2019, 6(7): 1801671

9. Qing Yuan, Yao Zhao, Pengju Cai, Zhesheng He, Fuping Gao, Jinsong Zhang, and Xueyun Gao, ACS Omega 2019, 4(9):14092–14099

10. Kean W. F., Hart L., Buchanan W. W. Auranofin. Br. J. Rheumatol. 1997, 36, 560–572

11. Xueyun Gao, 2018, US patent 9993562B2

12. J. M. Maderia, D. L. Gibson, W. F. Kean, A. Klegeris, Inflammopharmacology, 2012, 20, 297–296

13. Becke, A. D., Density-functional thermochemistry. III. The role of exact exchange. The Journal of Chemical Physics 1993, 98(7): 5648–5652.

14. Schwerdtfeger, P.; Dolg, M.; Schwarz, W. H. E.; Bowmaker, G. A.; Boyd, P. D. W., Relativistic effects in gold chemistry. I. Diatomic gold compounds. The Journal of Chemical Physics 1989, 91(3): 1762–1774.

15. Petersson, G. A.; Al - Laham, M. A., A complete basis set model chemistry. II. Open - shell systems and the total energies of the first - row atoms. The Journal of Chemical Physics 1991, 94(9): 6081–6090.

16. Marenich, A. V.; Cramer, C. J.; Truhlar, D. G., Universal Solvation Model Based on Solute Electron Density and on a Continuum Model of the Solvent Defined by the Bulk Dielectric Constant and Atomic Surface Tensions. The Journal of Physical Chemistry B 2009, 113(18):6378–6396.

17. Frisch, M. J.; Trucks, G. W.; Schlegel, H. B.; Scuseria, G. E.; Robb, M. A.; Cheeseman, J. R.; Scalmani, G.; Barone, V.; Mennucci, B.; Petersson, G. A.; Nakatsuji, H.; Caricato, M.; Li, X.; Hratchian, H. P.; Izmaylov, A. F.; Bloino, J.; Zheng, G.; Sonnenberg, J. L.; Hada, M.; Ehara, M.; Toyota, K.; Fukuda, R.; Hasegawa, J.; Ishida, M.; Nakajima, T.; Honda, Y.; Kitao, O.; Nakai, H.; Vreven, T.; Montgomery Jr., J. A.; Peralta, J. E.; Ogliaro, F.; Bearpark, M. J.; Heyd, J.; Brothers, E. N.; Kudin, K. N.; Staroverov, V. N.; Kobayashi, R.; Normand, J.; Raghavachari, K.; Rendell, A. P.; Burant, J. C.; Iyengar, S. S.; Tomasi, J.; Cossi, M.; Rega, N.; Millam, N. J.; Klene, M.; Knox, J. E.; Cross, J. B.; Bakken, V.; Adamo, C.; Jaramillo, J.; Gomperts, R.; Stratmann, R. E.; Yazyev, O.; Austin, A. J.; Cammi, R.; Pomelli, C.; Ochterski, J. W.; Martin, R. L.; Morokuma, K.; Zakrzewski, V. G.; Voth, G. A.; Salvador, P.; Dannenberg, J. J.; Dapprich, S.; Daniels, A. D.; Farkas, Ö.; Foresman, J. B.; Ortiz, J. V.; Cioslowski, J.; Fox, D. J. Gaussian 09, Gaussian, Inc.: Wallingford, CT, USA, 2009.

18. Phillips, J. C., Braun, R., Wang, W., Gumbart, J., Tajkhorshid, E., Villa, E., Chipot, C. Skeel, R. D. Kale, L. and Schulten, K.. Scalable molecular dynamics with NAMD. Journal of Computational Chemistry 2005, 26(16): 1781–1802.

19. Lee, J., Cheng, X., Swails, J. M., Yeom, M. S., Eastman, P. K., Lemkul, J. A., Wei, S., Buckner, J., Jeong, J. C., Qi, Y., Jo, S., Pande, V. S., Case, D. A., Brooks, C. L., MacKereall, Jr, A. D., Klauda, J. B. and Im, W. CHARMM-GUI input generator for NAMD, GROMACS, AMBER, OpenMM, and CHARMM/OpenMM simulations using the CHARMM36 additive force field. Journal of Chemical Theory and Computation, 2016, 12, 405–413.

